# MAGI2-AS3 modulates proliferation, apoptosis and glycolysis in acute lymphoblastic leukemia via miR-452-5p/FOXN3 axis

**DOI:** 10.1101/2021.07.02.450970

**Authors:** Xiao-Guang Chen, Bing-Hua Dou, Jin-Dou An, Song Feng, Na Liu, Guang-Yao Sheng

**Affiliations:** Department of Pediatric, The First Affiliated Hospital of Zhengzhou University, Zhengzhou 450052, Henan Province, P.R. China

**Keywords:** acute lymphoblastic leukemia, MAGI2-AS3, miR-452-5p, FOXN3, glycolysis

## Abstract

**Background:** Long non-coding RNA MAGI2 antisense RNA 3 (MAGI2-AS3) has been identified as a tumor suppressor in various cancers. Acute lymphoblastic leukemia (ALL) is a prevalent kind of leukemia among children. In this study, we aimed at evaluate the role of MAGI2-AS3 in ALL and its underlying mechanisms.

**Methods:** qPCR was adopted to determine MAGI2-AS3, miR-452-5p, and FOXN3 expression. The malignant properties of ALL cells were assessed by CCK8 assay and flow cytometry analysis. The glucose uptake, lactate production, and ATP level were measured to evaluate glycolysis. Western blotting was performed to detect PCNA, Bcl-2, Bax, and HK2 protein levels. The interaction between MAGI2-AS3/FOXN3 and miR-452-5p was validated by luciferase reporter assay. The *in vivo* growth of ALL cells was determined in xenograft model.

**Results:** MAGI2-AS3 was strikingly down-regulated in ALL samples and cells. Overexpression of MAGI2-AS3 restrained growth, glycolysis and triggered apoptosis of ALL cells. Mechanistically, MAGI2-AS3 could sponge miR-452-5p to up-regulate FOXN3. Silencing of FOXN3 abolished the anti-tumor effect of MAGI2-AS3. Finally, MAGI2-AS3 suppressed the *in vivo* growth of ALL cells via modulating miR-452-5p/FOXN3 axis.

**Conclusion:** Our findings demonstrate that MAGI2-AS3 delays the progression of ALL by regulating miR-452-5p/FOXN3 signaling pathway.

## Introduction

Acute lymphoblastic leukemia (ALL) is one of prevalent hematological malignant tumors among children, which accounts for about 75% of children’s acute leukemia[1]. According to immunophenotyping, ALL patients are generally divided into B-ALL or T-ALL. Although conventional chemotherapy has greatly improved the survival rates of ALL patients, the recurrent cases are hard to get remission[2]. In addition, adverse effects induced by conventional chemotherapy may increase the risk of developing a secondary tumor[3]. Hence, it is urgent to uncover the pathogenesis of ALL for providing novel therapeutic options.

Long non-coding RNAs (lncRNAs) are a kind of non-protein coding transcripts more than 200 nucleotides in length[4]. Growing evidence has revealed that lncRNAs are implicated in tumorigenesis and development of various malignancies[5, 6]. Previous publications have demonstrated that lncRNAs, served as tumor suppressor or promotor, play crucial roles in tumor occurrence, metastasis or recurrence[7, 8]. Besides, lncRNAs have also been confirmed to affect the tumorigenesis and progression of ALL[9]. MAGI2 antisense RNA 3 (MAGI2-AS3) has been documented to be down-regulated and play oncogenic roles in different types of malignancies, such as breast cancer, lung cancer, hepatocellular carcinoma, bladder cancer and so on[10]. Notably, a significant downregulation of MAGI2-AS3 has been found in ALL patient samples[11]. A recent study by Chen et al reported that MAGI2-AS3 exhibited a poor expression in acute myeloid leukaemia, which was responsible for self-renewal of leukaemia stem cells[12]. Yet, the biological functions of MAGI2-AS3 in ALL remains obscure.

MicroRNAs (miRNAs), comprised of 18-24 nucleotides, are non-coding RNAs possessing post-transcriptional modulatory potential of their target genes by binding to the 3′-untranslated region (3′-UTR)[13]. Mounting evidence has suggested that miRNAs engage in all kinds of biological processes, including cancer[14]. MiR-452-5p has been verified as tumor promoter in multiple tumors. For instance, increased expression of miR-452-5p facilitated colorectal cancer development via modulating PKN2/ERK/MAPK feedback loop[15]. In lung squamous cell carcinoma, miR-452-5p up-regulation remarkably correlated with tumor-node metastasis and exerted vital role in tumor progression via targeting CDKN1B[16]. However, whether miR-452-5p is involved in ALL development has not been revealed.

Forkhead box N3 (FOXN3), a member of forkhead box family, is involved various pathophysiological processes, including tumorigenesis[17]. As reported by previous researches, a decreased expression of FOXN3 is confirmed in a wide variety of tumors, including ALL[18, 19]. More importantly, bioinformatics analysis indicates that both MAGI2-AS3 and FOXN3 possess binding sites in miR-452-5p. Therefore, MAGI2-AS3 might modulate the progression of ALL through miR-452-5p/FOXN3 axis.

To validate the above hypothesis, this study probed into the regulatory function and molecular mechanisms of MAGI2-AS3 in ALL, which provided new insights into the pathogenesis of ALL.

## Materials and methods

### Clinical samples

Bone marrow specimens were collected from 25 ALL patients and 25 healthy cases after signing informed consent in The First Affiliated Hospital of Zhengzhou University. This study following the Declaration of Helsinki was approved by the Ethics Committee of The First Affiliated Hospital of Zhengzhou University.

### Cell culture

Human ALL cell lines Jurkat, Reh, CEM, and Molt4 were provided by the Cell Bank of the Chinese Academy of Sciences (Shanghai, China) and cultured in RPMI-1640 medium (Thermo Fisher, USA) containing 10% fetal bovine serum (Biological Industries, Israel). Peripheral blood mononuclear cells (PBMCs) were separated from the serum of normal subjects using density gradient centrifugation and maintained in RPMI-1640 medium with 10% fetal bovine serum.

### Cell transfection

The overexpression plasmid MAGI2-AS3 (OE-MAGI2-AS3), control vector, miR-452-5p mimics, mimics negative control (NC), short-hairpin RNA (shRNA) targeting FOXN3 (shFOXN3) and sh-NC were provided by GenePharma (Shanghai, China). Jurkat and Reh cells were transfected with corresponding segment using Lipofectamine 2000 (Thermo Fisher).

### Quantitative polymerase chain reaction (qPCR)

Total RNA was prepared using TRIzol agent (Thermo Fisher). Then, a Prime Script RT kit (Takara, Tokyo, Japan) was adopted for the generation of complementary DNA. The PCR amplifications were performed using the SYBR-Green PCR Master Mix (Applied Biosystems, USA). Evaluation of relative RNA levels was carried out with 2^−ΔΔCt^ method. Table 1 presents the primers of target genes.

**Table 1.**
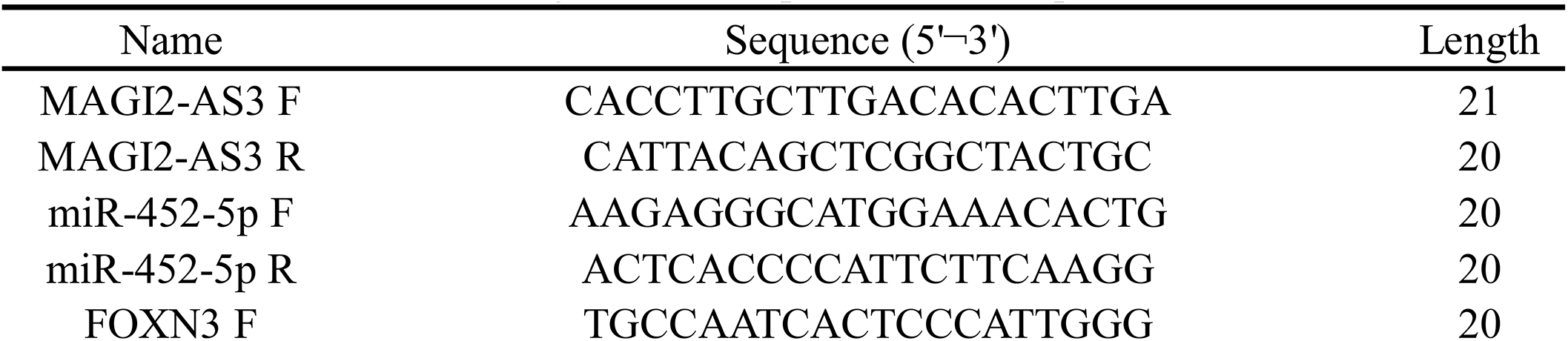
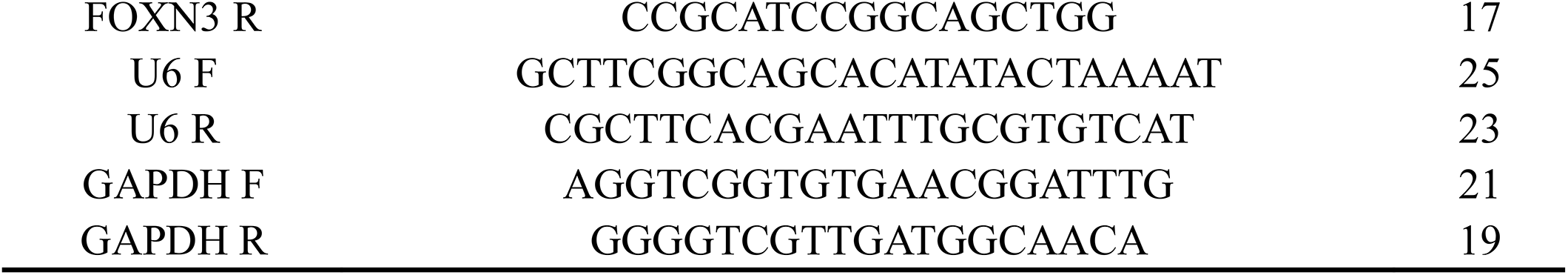
Oligonucleotide primer sets for qRT-PCR

### Cell counting kit-8 (CCK-8)

ALL cells seeded in 96-well plates were reacted with 10 μl of CCK-8 reagent (Dojindo, Japan) at 37°C at various time points. Four hours later, the absorbance intensity was detected at a wavelength of 450 nm on a microplate reader (Tecan, Switzerland).

### Flow cytometry

The apoptosis of ALL cells was assessed using a FITC Annexin V/Dead Cell Apoptosis Kit (Thermo Fisher). In short, the collected ALL cells were resuspended in binding buffer, followed by staining with FITC Annexin V (5 μL) and PI (1 μL) solution for 30 min away from light. Then the stained cells were detected on a flow cytometer (Millipore, USA).

### Detection of glucose uptake, lactate production and ATP level

To evaluate glycolysis of ALL cells, glucose uptake, lactate production and ATP level were determined using Glucose Uptake Assay Kit (Abcam, UK), Lactate Assay Kit II (Sigma-Aldrich, USA), and ATP Detection Kit (Solarbio, Beijing, China) following the protocols provided by the manufacturers.

### Western blotting

After lysis using Pierce IP lysis buffer (Thermo Fisher), total protein concentration was measured with a BCA kit (Thermo Fisher). Then, the protein samples were resolved by sodium dodecyl sulfate polyacrylamide gel electrophoresis and transferred to PVDF membranes. After sealing in 5% skim milk, probing with the primary antibodies against PCNA (ab92552, 1:1000, Abcam), Bcl-2 (ab32124, 1:1000, Abcam), Bax (ab32503, 1:1000, Abcam), HK2 (ab209847, 1:1000, Abcam), FOXN3 (#711585, 1:100, Thermo Fisher), GAPDH (#5174, 1:1000, CST, USA) at 4°C overnight. Subsequently, a secondary antibody (#A27036, 1:4000, Thermo Fisher) was applied. The bands were acquired using the enhanced chemiluminescence (ECL) reagent (Thermo Fisher).

### Dual-luciferase reporter assay

MAGI2-AS3 or FOXN3 3’UTR containing miR-452-5p wild-type (WT) or mutant (MUT) binding sites were inserted into psiCHECK vector (Promega, USA). 293T cells were transfected with the constructed luciferase plasmid together with miR-NC or miR-452-5p mimics using Lipofectamine 2000. The luciferase activities were assessed using a Dual-Lucy Assay Kit (Solarbio) at 48 h after the transfection.

### Animal experiment

BALB/c nude mice (18-22 g) were obtained from SLAC Laboratory Animal Co., Ltd (Shanghai, China). 1◻×◻10^7^ Jurkat cells stably transfected with OE-MAGI2-AS3 or vector were intraperitoneally implanted the nude mice (n=6 per group). Tumor length and width were measured every three days. Tumor volume was calculated as follow: ◻length ×◻width^2^/2 (mm^3^). Tumors were collected from mice 30 days later, weighed and subjected to subsequent experiments. All experimental protocols were approved by the Animal Ethics Committee of The First Affiliated Hospital of Zhengzhou University.

### TUNEL

To examine apoptosis in tumor tissues, the Colorimetric TUNEL Apoptosis Assay Kit (Beyotime) was adopted. Briefly, the tumor tissues were taken to paraffin imbedding and cut into 5-μm sections. After deparaffination and hydration, antigen retrieval was carried out using microwave. The sections were treated with Proteinase K (20 mg/mL) and then blocked in 3% H_2_O_2_ for 20 min. Thereafter, the reaction solution was added to the sections, followed by incubation at 37° C for 1 h. After reaction with streptavidin-HRP and DAB solution, the sections were observed under an optical microscope (Micro-shot Technology Co., Ltd, Guangzhou, China).

### Statistical analysis

Data are presented as mean± standard deviation (SD). GraphPad Prism 6.0 was adopted for statistical analysis using Student’s t test for two group or One-Way ANOVA for multiple group comparison. A p-value less than 0.05 was considered as statistically significant.

## Results

### Up-regulation of MAGI2-AS3 in ALL tissues and cells

First, we determined the differential expression of MAGI2-AS3 in bone marrow specimens from ALL patients and healthy volunteers. qPCR results revealed that MAGI2-AS3 was down-regulated in ALL bone marrow samples relative to normal controls (Fig. 1A). Consistently, a decreased expression of MAGI2-AS3 was validated in a series of ALL cells in comparison with PBMCs (Fig. 1B). Thus, MAGI2-AS3 might exert crucial regulatory roles in ALL progression.

**Figure 1.**
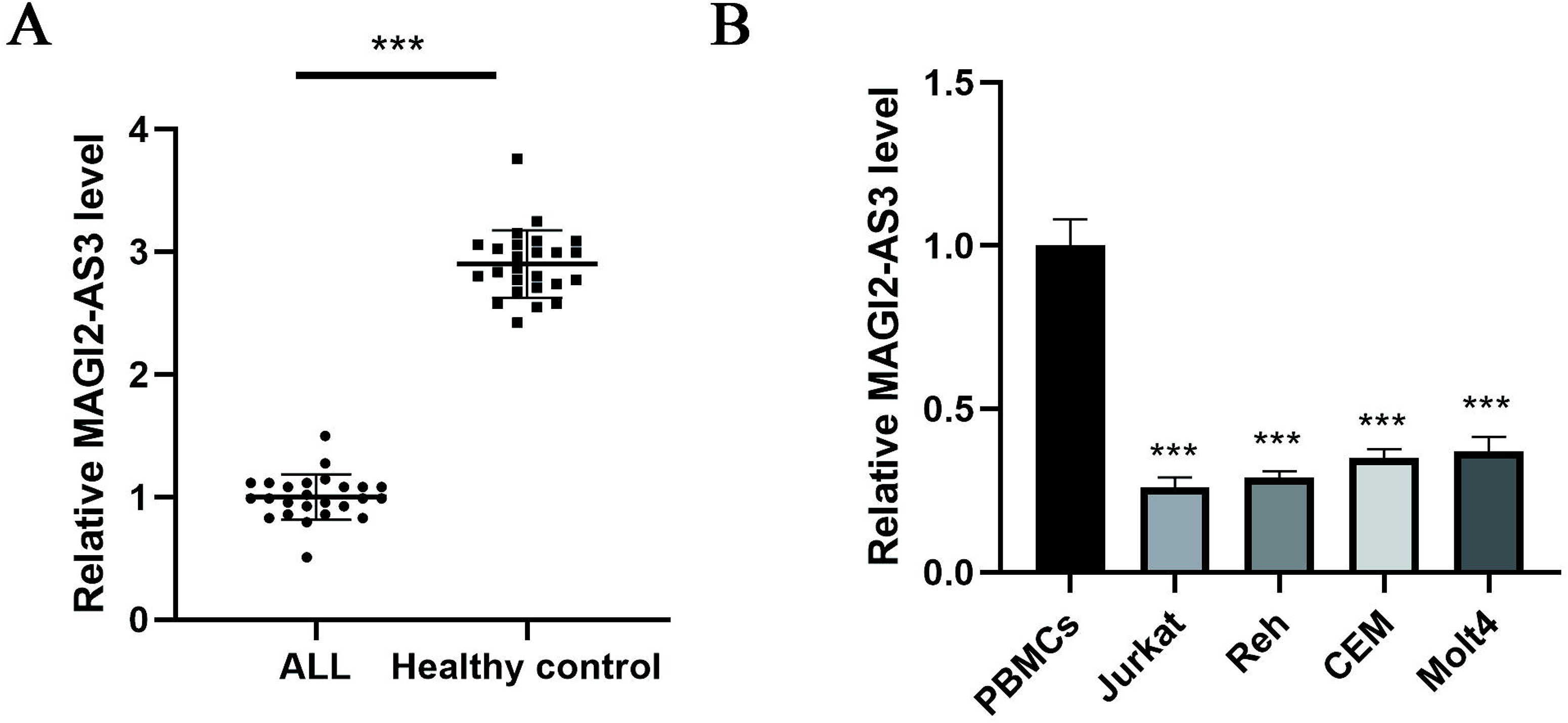
Aberrant high expression of MAGI2-AS3 in ALL tissues and cells. (A) qPCR for assessing MAGI2-AS3 level in the bone marrow samples of ALL and healthy controls. (B) The differential expression of MAGI2-AS3 among ALL cells and PBMCs was evaluated by qPCR. Data are expressed as mean ± SD. *** p < 0.001.

### Effect of MAGI2-AS3 on proliferation, apoptosis and glycolysis of ALL cells

To assess the regulation of MAGI2-AS3 in ALL, ALL cells were transfected with OE-MAGI2-AS3 plasmid to overexpress MAGI2-AS3. As verified by qPCR, OE-MAGI2-AS3 plasmid remarkably raised the expression of MAGI2-AS3 in Jurkat and Reh cells (Fig. 2A). Cell proliferation was determined by CCK8 assay, and we found that the proliferative ability was significantly reduced in MAGI2-AS3-overexpressed group (Fig. 2B). Flow cytometry was used to test the apoptosis of ALL cells. As illustrated in Fig. 2C, ectopic expression of MAGI2-AS3 evidently triggered apoptosis of Jurkat and Reh cells. Warburg effect in cancer cells has been well recognized, which is featured by an increase in glucose uptake and lactate production[20]. We demonstrated that overexpression of MAGI2-AS3 led to a striking decrease in glucose uptake, lactate production and ATP level in ALL cells (Fig. 2D-F). In addition, the protein levels of PCNA, Bcl-2, and HK2 were declined, but Bax level was elevated after overexpression of MAGI2-AS3 (Fig. 2G-I). These data implied that MAGI2-AS3 overexpression restrained proliferation, glycolysis and induced apoptosis of ALL cells.

**Figure 2.**
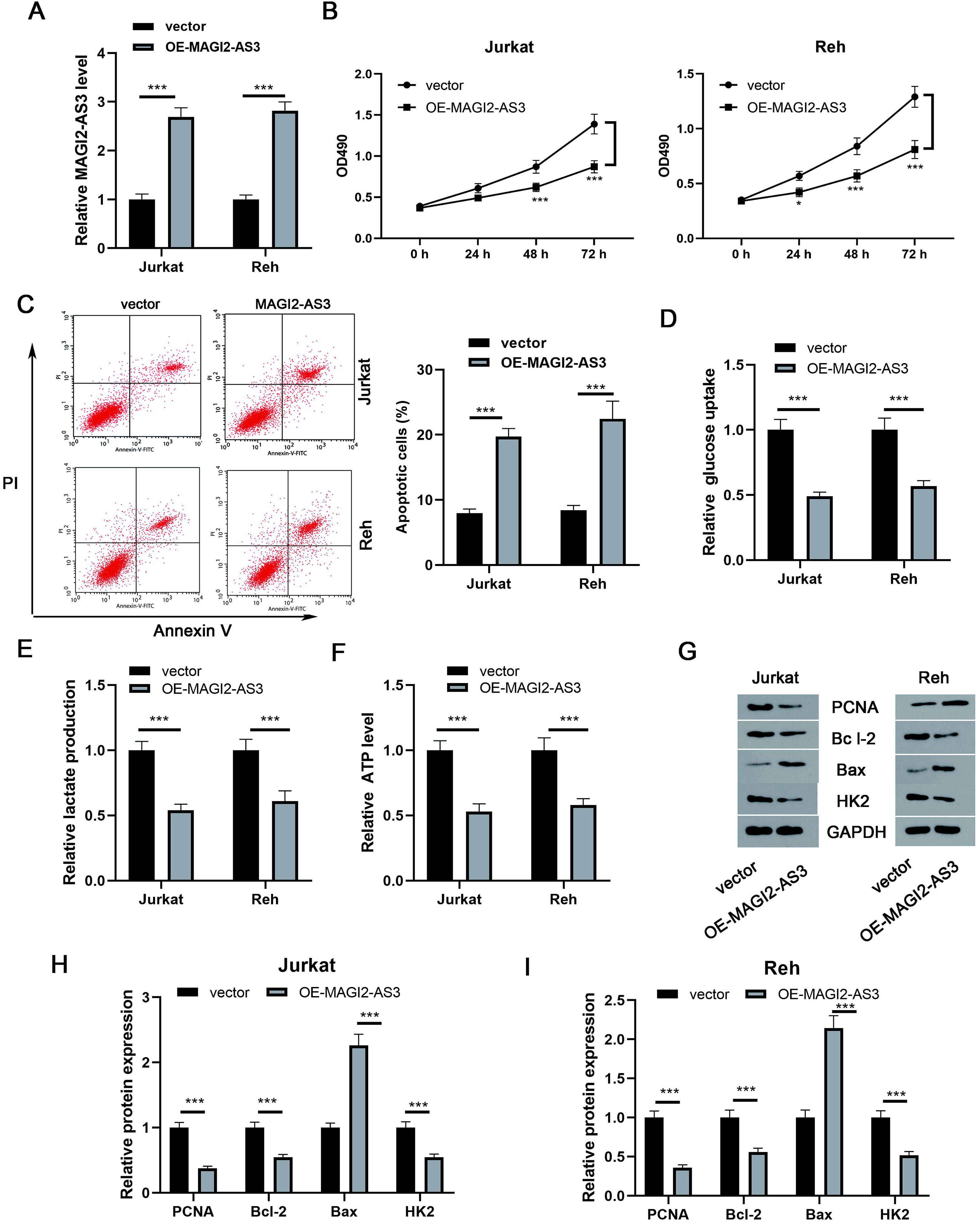
MAGI2-AS3 restrains proliferation, apoptosis inhibition and glycolysis of ALL cells. (A) MAGI2-AS3 overexpression efficiency in ALL cells was validated by qPCR. (B) Proliferation was assessed by CCK-8 after transfection with OE-MAGI2-AS3 plasmid. (C) Apoptosis was determined by flow cytometry. The glucose uptake (D), lactate production (E), and ATP level (F) were detected by commercial kits. (G-I) The protein levels of PCNA, Bcl-2, Bax, and HK2 were evaluated by Western blotting. Data are expressed as mean ± SD. * p < 0.05, *** p < 0.001.

### MAGI2-AS3 sponged miR-452-5p to enhance FOXN3 expression

To further investigate the underlying mechanisms of MAGI2-AS3, we focused on miR-452-5p. A reduction in miR-452-5p level was found in ALL cells after transfection with OE-MAGI2-AS3 (Fig. 3A). Additionally, miR-452-5p was up-regulated in the bone marrow samples of ALL cases (Fig. 3B). We also found a negative correlation between MAGI2-AS3 and miR-452-5p expression (Fig. 3C). Bioinformatics analysis demonstrated that MAGI2-AS3 could interact with miR-452-5p, and dual luciferase reporter assay revealed that miR-452-5p mimics obviously suppressed the luciferase activity of MAGI2-AS3-WT rather than MAGI2-AS3-MUT (Fig. 3D). Besides, both the mRNA and protein levels of FOXN3 in ALL cells were reduced by miR-452-5p overexpression (Fig. 3E-G). Accordingly, FOXN3 level was observably raised in ALL specimens (Fig. 3H), which was negative correlated with miR-452-5p level (Fig. 3I). Further bioinformatics analysis showed that FOXN3 was predicted as a target gene of miR-452-5p (Fig. 3J). Dual luciferase reporter assay suggested that miR-452-5p mimics evidently reduced the luciferase activity of FOXN3-3’UTR WT, however it did not affect that from FOXN3-3’UTR MUT group (Fig. 3K). qPCR and Western blotting analysis further demonstrated that overexpression of MAGI2-AS3 enhanced the mRNA and protein expression of FOXN3 in ALL cells, whereas miR-452-5p mimics transfection could abolish these changes (Fig. 3L-P). Collectively, MAGI2-AS3 up-regulated FOXN3 in ALL cells via sponging miR-452-5p.

**Figure 3.**
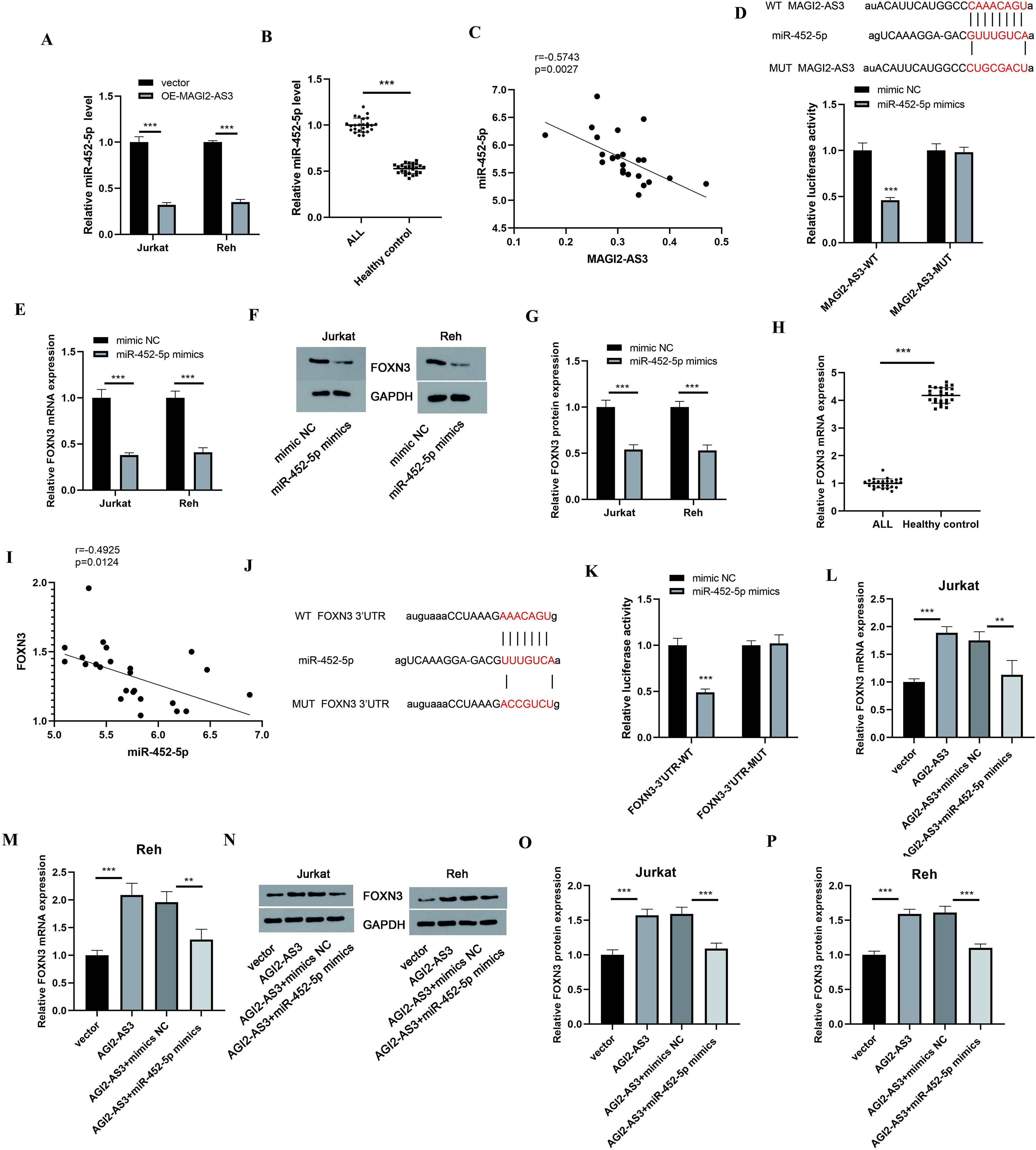
MAGI2-AS3 promotes FOXN3 expression via sponging miR-452-5p. (A) MiR-452-5p expression after transfection with OE-MAGI2-AS3 was measured by qPCR. (B) qPCR for detecting miR-452-5p level in ALL and healthy samples. (C) A negative correlation between MAGI2-AS3 and miR-452-5p level. (D) Dual-luciferase reporter assay for validating the direct binding between MAGI2-AS3 and miR-452-5p. FOXN3 level in miR-452-5p mimics-transfected ALL cells was determined by qPCR (E) and Western blotting (F-G). (H) FOXN3 mRNA expression in ALL and healthy samples was evaluated by qPCR. (I) FOXN3 negatively correlated with miR-452-5p level in ALL samples. (J) Putative binding sites between FOXN3 and miR-452-5p. (K) The interaction between FOXN3 and miR-452-5p was determined by dual-luciferase reporter assay. FOXN3 expression in ALL cells with multiple transfections was assessed by qPCR (L-M) and Western blotting (N-P). Data are expressed as mean ± SD. * p < 0.05, ** p < 0.01, *** p < 0.001.

### Silencing of FOXN3 reversed the anti-cancer effect of MAGI2-AS3

To determine the involvement of FOXN3 in MAGI2-AS3-mediated anti-cancer effects, the ALL cells were transfected with OE-MAGI2-AS3 together with or without shFOXN3. As detected by CCK8 assay, MAGI2-AS3 overexpression-induced proliferation inhibition of ALL cells could by restored by FOXN3 knockdown (Fig. 4A&B). In addition, the enhanced apoptosis of MAGI2-AS3-overexpressed ALL cells could be reversed by silencing of FOXN3 (Fig. 4C-E). Depletion of FOXN3 also recovered the decreased glucose uptake, lactate production, and ATP level induced by MAGI2-AS3 overexpression (Fig. 4F-K). Besides, transfection with OE-MAGI2-AS3 led to reduction in PCNA, Bcl-2, HK2 and elevation in Bax expression, which was partly counteracted by co-transfection with shFOXN3 (Fig. 4L-N). These findings suggested that MAGI2-AS3 delayed the progression of ALL via modulating FOXN3.

**Figure 4.**
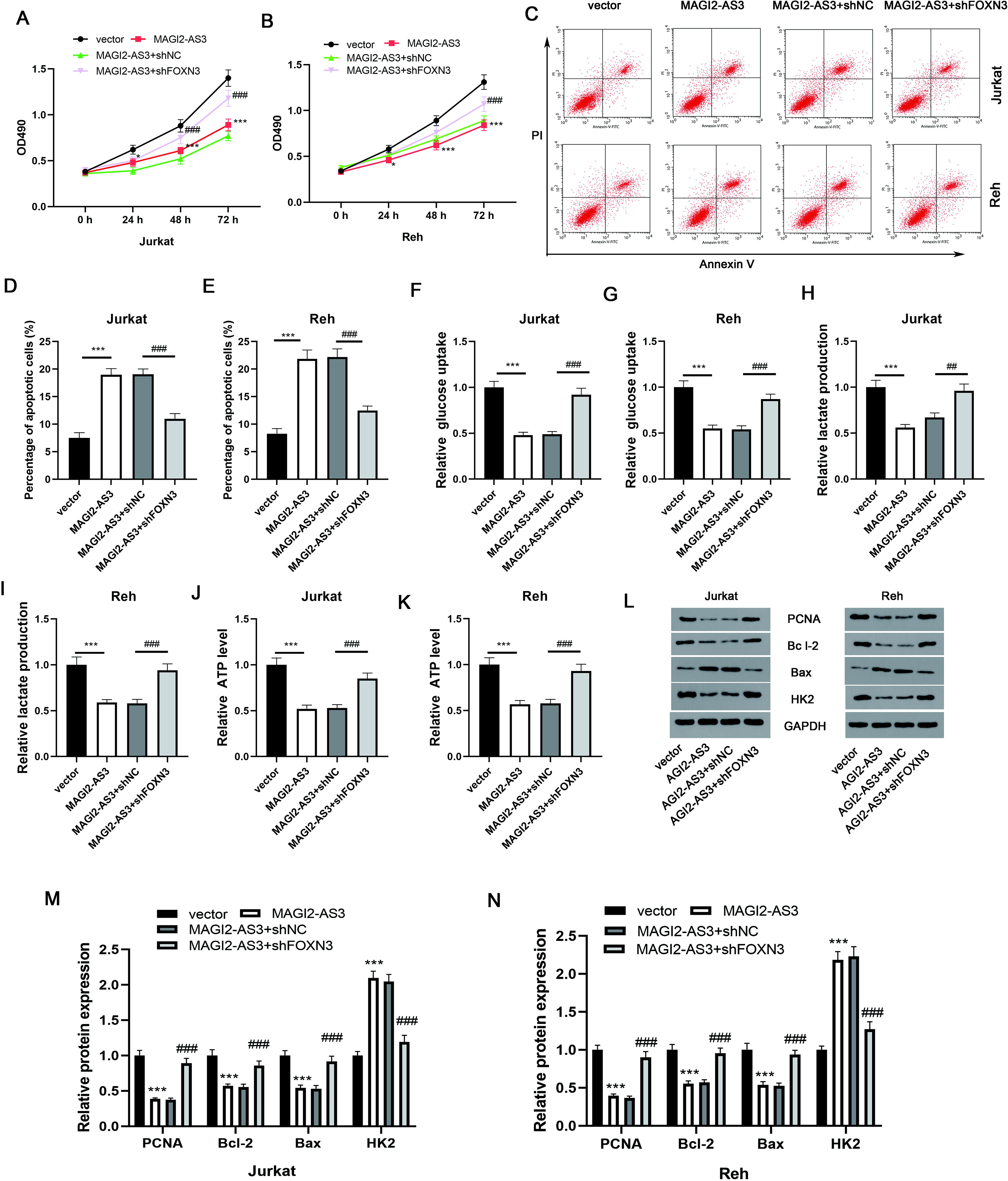
FOXN3 knockdown counteracts the anti-cancer effect of MAGI2-AS3. (A-B) CCK8 assay for determining proliferation of ALL cells from different groups. (C-E) Flow cytometry was adopted to assess apoptosis of ALL cells. (F-K) The glucose uptake, lactate production, and ATP levels were measured by commercial kits. (L-N) Western blotting for assessing PCNA, Bcl-2, Bax, and HK2 levels in ALL cells with various treatments. Data are expressed as mean ± SD. * p < 0.05, *** p < 0.001; ## p < 0.01, ### p < 0.001 versus the indicated group.

### Effect of MAGI2-AS3 on ALL tumor growth *in vivo*

To confirm the function of MAGI2-AS3 in tumorigenesis *in vivo*, the nude mice were inoculated with Jurkat cells stably transfected with OE-MAGI2-AS3 or vector. As presented in Fig. 5A-C, the tumor volume and weight were distinctly reduced by overexpression of MAGI2-AS3. Moreover, TUNEL assay revealed that apoptosis in tumor tissues was enhanced in response to MAGI2-AS3 overexpressing (Fig. 5D). Furthermore, the expression of MAGI2-AS3 and FOXN3 was elevated, whereas the miR-452-5p level was declined in the tumor tissues generated from MAGI2-AS3-overexpressed cells (Fig. 5E). Taken together, MAGI2-AS3 inhibited the growth of ALL xenografts via regulating miR-452-5p/FOXN3 axis.

**Figure 5.**
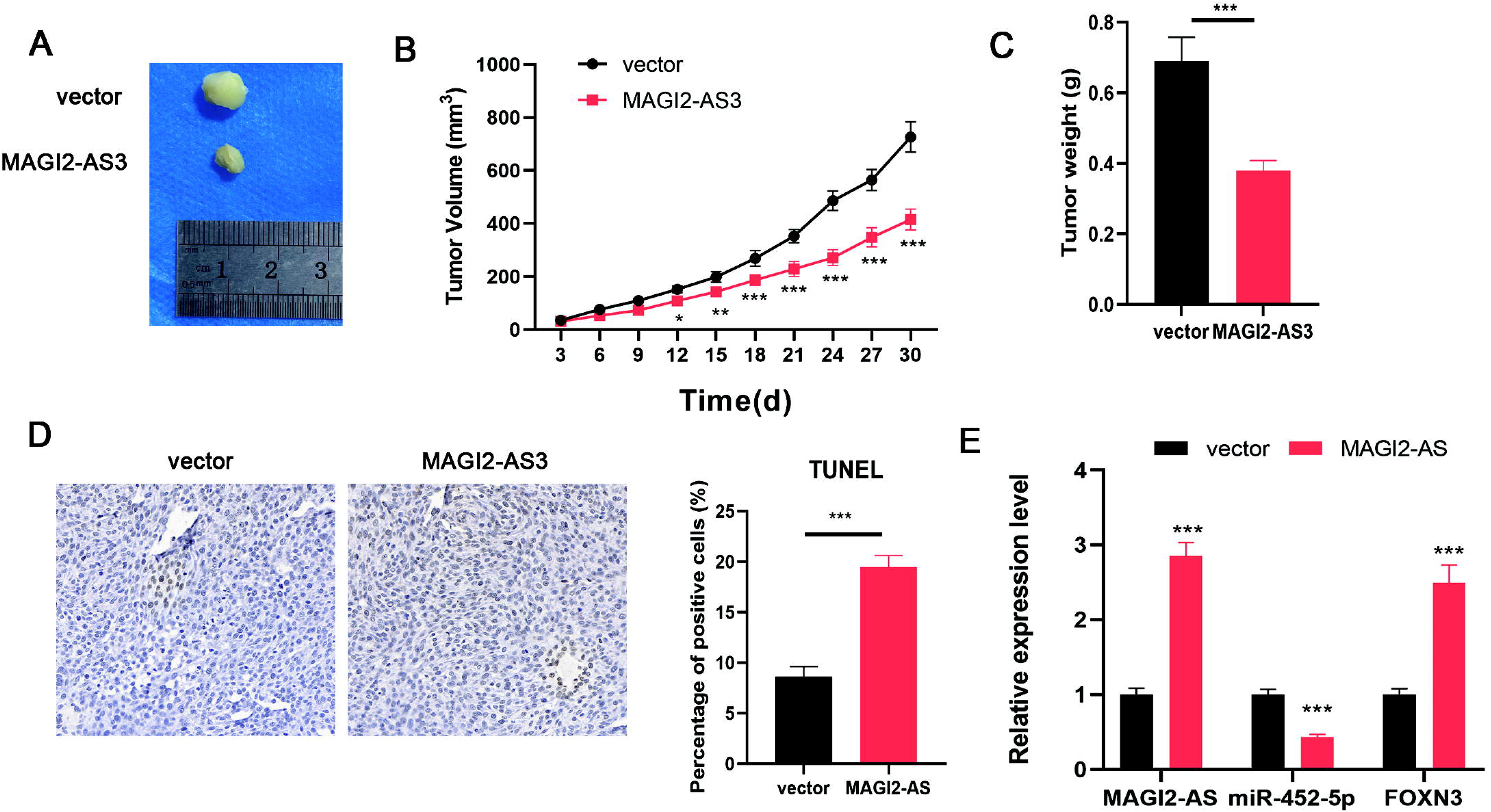
MAGI2-AS3 silencing inhibits ALL tumor growth *in vivo.* (A) Representative image of xenograft was presented. Tumor volume (B) and weight (C) from different groups were recorded. (D) TUNEL was used to determine apoptosis in tumor tissues. (E) Expression of MAGI2-AS3, FOXN3, and miR-452-5p in tumor tissues was detected by qPCR. Data are expressed as mean ± SD. * p < 0.05, ** p < 0.01, *** p < 0.001.

## Discussion

Over the past several decades, great improvement has been made in understanding the pathogenesis of ALL. However, the detailed mechanisms underlying the malignant progression and recurrence of ALL cells remain not entirely clear. This study focused on a novel modulatory pathway MAGI2-AS3/miR-452-5p/FOXN3 during ALL development. We demonstrated that MAGI2-AS3 and FOXN3 were up-regulated, and miR-345-5p was down-regulated in ALL specimens and cells. MAGI2-AS3 sequestered miR-345-5p to facilitate FOXN3 expression, which contributed to ALL cell growth, apoptosis inhibition and glycolysis.

MAGI2-AS3, located on chromosome 7q21.11, is reported to modulate the progression of a series of malignancies[21]. MAGI2-AS3 affects the malignant properties of tumors through various mechanisms, such as ceRNAs, epigenomic or transcriptional regulations. For instance, MAGI2-AS3 could favor colorectal cancer cell growth and metastasis by sponging miR-3163[22]. MAGI2-AS3 increased MAGI2 level via suppressing DNA demethylation of MAGI2, delaying proliferation and migration of breast cancer cells[23]. Additionally, MAGI2-AS3 transcriptionally down-regulated HOXB7, which slowed the development of esophageal cancer[24]. Recently, the carcinogenic role of MAGI2-AS3 in acute myeloid leukemia has been identified[12]. According to our data, MAGI2-AS3 was strikingly lowly expressed in ALL clinical samples and cells. Furthermore, enforced expression of MAGI2-AS3 suppressed proliferation and apoptosis inhibition of ALL cells. Thus, our study was the first to reveal that MAGI2-AS3 functioned as a tumor suppressor lncRNA during ALL development.

Glucose metabolic reprogramming has been considered as a striking hallmark of tumors[25]. Being different from healthy cells, tumor cells are prone to metabolize glucose by glycolysis, leading to enhanced glucose uptake and lactate production[26]. Glycolysis confers the malignant phenotypes as well as drug resistance of ALL cells[27]. HK2 has been recognized as a specific enzyme catalyzing glycolysis process[28]. In this study, we demonstrated that MAGI2-AS3 overexpression restrained glucose uptake, lactate production, ATP level, as well as reduced HK2 expression in ALL cells. Therefore, repression of glycolysis was involved in the anti-cancer effect of MAGI2-AS3 in ALL.

To further probe the regulatory mechanisms of MAGI2-AS3 in ALL, we focused on the ceRNA hypothesis that cytoplasmic lncRNA adsorbs miRNA to release the expression of miRNA target gene. miR-452-5p is a crucial member of miRNA, and its involvement in cancers has been reported. The interaction between miR-452-5p and lncRNAs has been identified by various studies. Song et al showed that lncRNA SOX2-OT facilitated the tumorigenesis of prostate cancer by suppressing miR-452-5p expression[29]. Zhu et al indicated that LINC00052 reversed the malignant behavior of hepatocellular carcinoma cells via sponging miR-452-5p[30]. Our data showed that miR-452-5p was aberrantly highly expressed in ALL, which was negatively correlated with MAGI2-AS3 level. Moreover, the direct binding relationship between MAGI2-AS3 and miR-452-5p was validated. These data indicated that MAGI2-AS3 down-regulation resulted in abnormal higher miR-452-5p expression, which contributed to ALL progression.

As comprehensively reported by previous researches, FOXN3 can function as a tumor suppressor gene in multiple tumors. FOXN3 could repress the growth and metastasis of colon cancer cells via inactivating β-catenin/TCF pathway[31]. More importantly, a decreased transcript level of FOXN3 was verified in ALL cells[19]. Consistent with previous observations, we demonstrated that FOXN3 down-regulated in ALL patients and cells. In addition, FOXN3 was validated to be a target of miR-452-5p. FOXN3 level could be enhanced by MAGI2-AS3, while reduced by miR-452-5p. The inhibition of MAGI2-AS3 in ALL cell growth, apoptosis suppression and glycolysis could be reversed by silencing of FOXN3. Collectively, MAGI2-AS3/miR-452-5p/FOXN3 axis was implicated in the pathogenesis of ALL. To sum up, we recognized MAGI2-AS3 as a tumor suppressor in ALL. MAGI2-AS3 overexpression repressed growth, glycolysis and induced apoptosis of ALL cells via sequestering miR-452-5p to elevate FOXN3 level. This work helps to further uncover the pathological mechanisms of ALL and identifies MAGI2-AS3 as a potential diagnostic and therapeutic target.

## Acknowledgements

None.

## Author contributions

XGC and GYS designed research; XGC, BHD and JDA developed the theory and performed the study; XGC, BHD, SF and NL performed the data analysis. GYS reviewed the data and figure; XGC and GYS wrote the paper. All authors read and approved the final manuscript.

## References

1 Siegel, R., Naishadham, D., Jemal, A. 2013 Cancer statistics, 2013. CA Cancer J Clin. 63, 11–30. (10.3322/caac.21166)

2 Li, C., Zhao, T., Nie, L., Zou, Y., Zhang, Q. 2020 MicroRNA-223 decreases cell proliferation, migration, invasion, and enhances cell apoptosis in childhood acute lymphoblastic leukemia via targeting Forkhead box O 1. Biosci Rep. 40, (BSR20200485 [pii] 10.1042/BSR20200485226491 [pii])

3 Robison, L. L., Bhatia, S. 2003 Late-effects among survivors of leukaemia and lymphoma during childhood and adolescence. Br J Haematol. 122, 345–359. (4499 [pii] 10.1046/j.1365-2141.2003.04499.x)

4 Gugnoni, M., Ciarrocchi, A. 2019 Long Noncoding RNA and Epithelial Mesenchymal Transition in Cancer. Int J Mol Sci. 20, (E1924 [pii] 10.3390/ijms20081924 ijms20081924 [pii])

5 Hosseini, E. S., Meryet-Figuiere, M., Sabzalipoor, H., Kashani, H. H., Nikzad, H., Asemi, Z. 2017 Dysregulated expression of long noncoding RNAs in gynecologic cancers. Mol Cancer. 16, 107. (10.1186/s12943-017-0671-2 10.1186/s12943-017-0671-2 [pii])

6 Farooqi, A. A., Attar, R., Yulaevna, I. M., Berardi, R. 2021 Interaction of long non-coding RNAs and circular RNAs with microRNAs for the regulation of immunological responses in human cancers. Semin Cell Dev Biol. (S1084-9521(21)00138-5 [pii] 10.1016/j.semcdb.2021.05.029)

7 Xu, K., Zhang, Z., Qian, J., Wang, S., Yin, S., Xie, H., Zhou, L., Zheng, S. 2019 LncRNA FOXD2-AS1 plays an oncogenic role in hepatocellular carcinoma through epigenetically silencing CDKN1B(p27) via EZH2. Exp Cell Res. 380, 198–204. (S0014-4827(19)30190-9 [pii] 10.1016/j.yexcr.2019.04.016)

8 Gu, X., Chu, Q., Zheng, Q., Wang, J., Zhu, H. 2021 The dual functions of the long noncoding RNA CASC15 in malignancy. Biomed Pharmacother. 135, 111212. (S0753-3322(20)31405-0 [pii] 10.1016/j.biopha.2020.111212)

9 Bahari, G., Hashemi, M., Naderi, M., Sadeghi-Bojd, S., Taheri, M. 2018 Long non-coding RNA PAX8-AS1 polymorphisms increase the risk of childhood acute lymphoblastic leukemia. Biomed Rep. 8, 184–190. (10.3892/br.2017.1028BR-0-0-1028 [pii])

10 Xue, C., Li, G., Lu, J., Luo, J., Jia, J. 2021 Novel insights for lncRNA MAGI2-AS3 in solid tumors. Biomed Pharmacother. 137, 111429. (10.1016/j.biopha.2021.111429)

11 Garitano-Trojaola, A., Jose-Eneriz, E. S., Ezponda, T., Unfried, J. P., Carrasco-Leon, A., Razquin, N., Barriocanal, M., Vilas-Zornoza, A., Sangro, B., Segura, V., et al. 2018 Deregulation of linc-PINT in acute lymphoblastic leukemia is implicated in abnormal proliferation of leukemic cells. Oncotarget. 9, 12842–12852. (10.18632/oncotarget.24401)

12 Chen, L., Fan, X., Zhu, J., Chen, X., Liu, Y., Zhou, H. 2020 LncRNA MAGI2-AS3 inhibits the self-renewal of leukaemic stem cells by promoting TET2-dependent DNA demethylation of the LRIG1 promoter in acute myeloid leukaemia. RNA Biol. 17, 784–793. (10.1080/15476286.2020.1726637)

13 Liu, X., Cai, H., Sheng, W., Huang, H., Long, Z., Wang, Y. 2018 microRNAs expression profile related with response to preoperative radiochemotherapy in patients with locally advanced gastric cancer. BMC Cancer. 18, 1048. (10.1186/s12885-018-4967-4)

14 Inoue, J., Inazawa, J. 2021 Cancer-associated miRNAs and their therapeutic potential. J Hum Genet. (10.1038/s10038-021-00938-6)

15 Lin, X., Han, L., Gu, C., Lai, Y., Lai, Q., Li, Q., He, C., Meng, Y., Pan, L., Liu, S., et al. 2021 MiR-452-5p promotes colorectal cancer progression by regulating an ERK/MAPK positive feedback loop. Aging (Albany NY). 13, 7608–7626. (10.18632/aging.202657)

16 Gan, X. N., Gan, T. Q., He, R. Q., Luo, J., Tang, R. X., Wang, H. L., Zhou, H., Qing, H., Ma, J., Hu, X. H., et al. 2018 Clinical significance of high expression of miR-452-5p in lung squamous cell carcinoma. Oncol Lett. 15, 6418–6430. (10.3892/ol.2018.8088)

17 Zhang, J., Wang, Y., Mo, W., Zhang, R., Li, Y. 2020 The clinical and prognostic significance of FOXN3 downregulation in acute myeloid leukaemia. Int J Lab Hematol. 42, 270–276. (10.1111/ijlh.13162)

18 Kong, X., Zhai, J., Yan, C., Song, Y., Wang, J., Bai, X., Brown, J. A. L., Fang, Y. 2019 Recent Advances in Understanding FOXN3 in Breast Cancer, and Other Malignancies. Front Oncol. 9, 234. (10.3389/fonc.2019.00234)

19 Nagel, S., Pommerenke, C., Meyer, C., Kaufmann, M., MacLeod, R. A. F., Drexler, H. G. 2017 Identification of a tumor suppressor network in T-cell leukemia. Leuk Lymphoma. 58, 2196–2207. (10.1080/10428194.2017.1283029)

20 Li, J., Huang, Q., Long, X., Guo, X., Sun, X., Jin, X., Li, Z., Ren, T., Yuan, P., Huang, X., et al. 2017 Mitochondrial elongation-mediated glucose metabolism reprogramming is essential for tumour cell survival during energy stress. Oncogene. 36, 4901–4912. (10.1038/onc.2017.98)

21 Kai-Xin, L., Cheng, C., Rui, L., Zheng-Wei, S., Wen-Wen, T., Peng, X. 2021 Roles of lncRNA MAGI2-AS3 in human cancers. Biomed Pharmacother. 141, 111812. (10.1016/j.biopha.2021.111812)

22 Ren, H., Li, Z., Tang, Z., Li, J., Lang, X. 2020 Long noncoding MAGI2-AS3 promotes colorectal cancer progression through regulating miR-3163/TMEM106B axis. J Cell Physiol. 235, 4824–4833. (10.1002/jcp.29360)

23 Xu, X., Yuan, X., Ni, J., Guo, J., Gao, Y., Yin, W., Li, F., Wei, L., Zhang, J. 2021 MAGI2-AS3 inhibits breast cancer by downregulating DNA methylation of MAGI2. J Cell Physiol. 236, 1116–1130. (10.1002/jcp.29922)

24 Cheng, W., Shi, X., Lin, M., Yao, Q., Ma, J., Li, J. 2020 LncRNA MAGI2-AS3 Overexpression Sensitizes Esophageal Cancer Cells to Irradiation Through Down-Regulation of HOXB7 via EZH2. Front Cell Dev Biol. 8, 552822. (10.3389/fcell.2020.552822)

25 Pavlova, N. N., Thompson, C. B. 2016 The Emerging Hallmarks of Cancer Metabolism. Cell Metab. 23, 27–47. (10.1016/j.cmet.2015.12.006)

26 Liberti, M. V., Locasale, J. W. 2016 Correction to: ‘The Warburg Effect: How Does it Benefit Cancer Cells?’: [Trends in Biochemical Sciences, 41 (2016) 211]. Trends Biochem Sci. 41, 287. (10.1016/j.tibs.2016.01.004)

27 Hulleman, E., Kazemier, K. M., Holleman, A., VanderWeele, D. J., Rudin, C. M., Broekhuis, M. J., Evans, W. E., Pieters, R., Den Boer, M. L. 2009 Inhibition of glycolysis modulates prednisolone resistance in acute lymphoblastic leukemia cells. Blood. 113, 2014–2021. (10.1182/blood-2008-05-157842)

28 Liberti, M. V., Locasale, J. W. 2016 The Warburg Effect: How Does it Benefit Cancer Cells? Trends Biochem Sci. 41, 211–218. (10.1016/j.tibs.2015.12.001)

29 Song, X., Wang, H., Wu, J., Sun, Y. 2020 Long Noncoding RNA SOX2-OT Knockdown Inhibits Proliferation and Metastasis of Prostate Cancer Cells Through Modulating the miR-452-5p/HMGB3 Axis and Inactivating Wnt/beta-Catenin Pathway. Cancer Biother Radiopharm. 35, 682–695. (10.1089/cbr.2019.3479)

29 Zhu, L., Yang, N., Chen, J., Zeng, T., Yan, S., Liu, Y., Yu, G., Chen, Q., Du, G., Pan, W., et al. 2017 LINC00052 upregulates EPB41L3 to inhibit migration and invasion of hepatocellular carcinoma by binding miR-452-5p. Oncotarget. 8, 63724–63737. (10.18632/oncotarget.18892)

29 Dai, Y., Wang, M., Wu, H., Xiao, M., Liu, H., Zhang, D. 2017 Loss of FOXN3 in colon cancer activates beta-catenin/TCF signaling and promotes the growth and migration of cancer cells. Oncotarget. 8, 9783–9793. (10.18632/oncotarget.14189)

